# RTF1 mediates epigenetic control of Th17 cell differentiation via H2B monoubiquitination

**DOI:** 10.1101/2024.11.08.622668

**Authors:** Carolina Galan, Guangqing Lu, Richard Gill, Dun Li, Yifang Liu, Jun R. Huh, Saiyu Hang

## Abstract

A gene encoding the transcription factor RTF1 has been associated with an increased risk of ulcerative colitis (UC). In this study, we investigated its function in modulating T cells expressing interleukin-17A (Th17 cells), a cardinal cell type promoting intestinal inflammation. Our results indicate that *Rtf1* deficiency disrupts the differentiation of Th17 cells, while leaving regulatory T cells (Treg) unaffected. Mechanistically, RTF1 facilitates histone H2B monoubiquitination (H2Bub1), which requires its histone modification domain (HMD), for supporting Th17 cell function. Impaired Th17 differentiation was also observed in cells lacking the H2Bub1 E3 ligase subunit RNF40, an enzyme known to physically interact with RTF1. Thus, our study underscores the essential role of RTF1 in H2Bub1-mediated epigenetic regulation of Th17 cell differentiation. Understanding this process will likely provide valuable insights into addressing Th17-associated inflammatory disorders.

**Key Points:**

- RTF1 is crucial for Th17 cell differentiation via histone H2B monoubiquitination
- RTF1 deficiency impairs Th17 differentiation, similar to H2Bub1 ligase RNF40 deficiency
- RTF1 and H2Bub1 colocalize on genes regulating Th17 cells, indicating an epigenetic role

## Introduction

Epigenetic modifications play a critical role in transcriptional regulation and the differentiation of many cell types, including immune cells (Wei et al., 2009). Ample evidence suggests that nucleosome positioning and histone modifications, such as acetylation and methylation, correlate with gene transcription in distinct T cell subsets (Barski et al., 2007; Roh et al., 2005; Schones et al., 2008). For example, in Th1 cells, interferon-γ (IFN-γ) gene expression is facilitated by active histone marks, such as H3K4 trimethylation and H3K9 acetylation, while in Th2 cells, *Gata3* and *Il-4* loci were found with enhanced H3K27 acetylation and H3K4 methylation, bolstering cytokine production (Aune et al., 2009; Wei et al., 2009, 2011). The histone modification landscape in Th17 cells is also dynamically regulated, with H3K4 and H3K27 trimethylation associated with pro-inflammatory gene activation (Cribbs et al., 2020; Wei et al., 2009). Conversely, Treg cells exhibit distinct histone modifications at the *Foxp3* locus, where permissive marks, like H3K4me3, and repressive marks, such as H3K27me3, establish a balance between FOXP3 expression and stability (Kitagawa et al., 2017; Wei et al., 2009). In addition to the methylation at the histone subunits, monoubiquitination of histone H2B at lysine residue 120 (H2Bub1) has emerged as a key posttranslational modification associated with actively transcribed genes (Jung et al., 2012; Fuchs et al., 2014; Bourbousse et al., 2012). However, the precise impact of H2Bub1 on CD4^+^ T cell subset differentiation remains elusive. Several studies performed in model organisms have identified a role for the transcription factor RTF1 (Restores TBP function 1) in histone modification. RTF1 was initially identified as a suppressor of a TATA-binding protein (TBP) mutant in yeast (Stolinski et al., 1997). Subsequent genetic and biochemical studies in yeast have shown that RTF1 functions as a component of the polymerase-associated factor 1 (Paf1) complex (PAF1C), which consists of Paf1, Ctr9, Leo1, and Cdc73 (Mueller and Jaehning, 2002; Shi et al., 1997; Squazzo et al., 2002). The primary role of PAF1C is to facilitate transcription elongation (Kim et al., 2010; Rondón et al., 2004). Recent studies in mice and humans, however, suggest RTF1 is not stably associated with the PAF1C (Cao et al., 2015; Parnas et al., 2015). In yeast, RTF1 is also known to bind the chromatin remodeler CHD1 (Simic et al., 2003) and is required for the monoubiquitination of histone H2B (H2Bub1) (Ng et al., 2003; Wood et al., 2003; Xiao et al., 2005) and histone methylation (Warner et al., 2007). *Drosophila* RTF1 also functions in histone methylation, gene expression, and Notch signaling (Tenney et al., 2006).

In this study, we identified RTF1 from a reporter cell-based genetic screen and analyzed the genetic association of *RTF1* variants with ulcerative colitis. Using various genetic knockout (KO) models, we probed the modulatory role of RTF1 in the differentiation and function of Th17 cells, one of the major cell types promoting gut inflammatory pathologies (Brand, 2009; Blaschitz and Raffatellu, 2010; Gálvez, 2014; Leppkes et al., 2009). Our results revealed that RTF1 deficiency markedly disrupts the expression of Th17-specific genes. Notably, the deficiency of RTF1 does not impact the expression of *Rorc* (RAR-related orphan receptor C, the gene that encodes RAR-related orphan receptor gamma), a pivotal transcription factor for Th17 development. Moreover, RTF1 is critical for the maintenance of H2Bub1, which promotes Th17 differentiation. Intriguingly, mice deficient in the H2B ubiquitin ligase RNF40 also had impaired Th17 cell differentiation. Functionally, the histone modification domain (HMD) of RTF1 is responsible for H2Bub1. Indeed, RTF1’s HMD mutant form no longer supports H2Bub1 and Th17 differentiation. Finally, RTF1 and H2Bub1 colocalize on multiple genes known to regulate Th17. Thus, RTF1-dependent H2Bub1 represents a novel epigenetic activation mechanism contributing to Th17 cell development and function.

## Materials and methods

### Colocalization Analysis of UC GWAS and GTEx eQTL data

To evaluate the effect of UC GWAS loci on *RTF1* expression, we used coloc (Giambartolomei et al., 2014; Wang et al., 2020) to perform colocalization analysis of UC summary association statistics from Liu et al. 2015 (Liu et al., 2015) and the GTEx v8 eQTL catalog. We ran coloc to test whether the disease-and expression-associated signals at a given locus were colocalized, assuming there is at most one causal variant for each pair of disease and expression traits. Coloc uses summary statistics from trait pairs as Bayesian priors to calculate approximate Bayes factors representing the posterior probability of five hypotheses: no association with either trait (H0), association with trait 1 but not trait 2 (H1), association with trait 2 but not trait 1 (H2), association with both traits at correlated but distinct causal variants (H3), or colocalization: association with both traits at a shared causal variant (H4). To visually confirm significantly colocalizing loci we generated and examined using locuszoom plots with stacked disease-and gene expression-associated genetic loci (Pruim et al., 2010).

### dsRNA screening in *Drosophila* S2 cell reporter system

RORγt and other reporter cell lines in *Drosophila* S2 cells were previously described(Huh et al., 2011). RORγ/γt-LBD-Gal4-DBD driven luciferase reporter assays were performed in conjunction with a high-throughput loss of function screen using a dsRNA library. *Drosophila* dsRNA screen was performed in the *Drosophila* RNAi screening center (DRSC) at Harvard Medical School according to a protocol provided by the core, described elsewhere (Flockhart et al., 2006).

### Mice

*Rtf1* floxed mice were derived from *Rtf1^tm1a(KOMP)Wtsi^* embryonic stem cells (Project ID: CSD31048) sourced from the KOMP, which were microinjected into blastocyst for generating mice and subsequently crossed with FLP transgenic animals. The resulting offspring were then bred with *Cd4-Cre* (Jax # 017336), UBC-Cre-ERT2 (Jax # 007001), and *Il17a-Cre* (Jax # 016879). C57BL/6 and *Rag1-KO* mice were obtained from Jackson laboratory or Taconic Biosciences. *Rnf40-flox* mice were generated through homologous recombination in mouse ES cells, incorporating loxP sites flanking *Rnf40* exon 3-5. Rosa26-ERT2 mice were procured from Taconic Biosciences. All mice were maintained in specific pathogen-free facilities, and all animal experiments were conducted in accordance with an IACUC-approved protocol at Harvard Medical School, and in Genentech.

### In vitro T cell culture and flow cytometric analyses

Naive CD4^+^ T cells (CD4^+^CD62L^hi^CD44^low^) were MACS purified (Miltenyi Biotech) and sorted from the spleens and lymph nodes of mice. The cells were cultured and stimulated with anti-CD3 (clone 145-2C11; ThermoFisher, 5 µg/mL) and anti-CD28 (clone 37.51; ThermoFisher, 5 µg/mL) under different cytokine cocktails. The T cells were cultured with hIL-2 (Peprotech # 200-02, 100 U/mL) for Th0 differentiation, mIL-6 (Peprotech # 216-16, 20 ng/mL) and hTGFβ1 (Peprotech # 100-21, 0.5 ng/mL) for Th17 differentiation, hIL-2 (100 U/mL) and hTGFβ1 (1 ng/mL) for iTreg differentiation. DMSO or 4-hydroxytamoxifen (1 nM) was added from the beginning of the culture.

After three days of culture, cells were treated with 50 ng/mL PMA (Phorbol 12-myristate 13-acetate, Sigma) and 1 µM ionomycin (Sigma) along with GolgiPlug (BD) for 4 hours to induce cytokine production. Following stimulation, cells were labeled with cell surface marker antibodies and LIVE/DEAD fixable dye. They were subsequently fixed and permeabilized using the FOXP3/Transcription factor staining kit (eBiosciences), following the manufacturer’s guidelines, and then stained with antibodies specific to cytokine and/or transcription factors. The staining antibodies were purchased from eBioscience or Biolegend. All acquisitions were performed with a BD LSR II flow cytometer and were analyzed with FlowJo software v9 (TreeStar).

### Immunoblotting

Sorting purified or cells cultured in vitro were harvested, and total protein extracts were solubilized in SDS sample buffer. The extracts were separated on 12% SDS-PAGE gels and transferred to polyvinylidene difluoride membrane (Millipore). The membranes were probed with specific antibodies targeting RTF1 (Sigma), β-actin (Cell Signaling), H2Bub1(clone D11, Cell Signaling Technology), and H2B (clone D2H6, Cell Signaling Technology). Visualization was achieved using the ECL HRP Substrate (ThermoFisher).

### Immunofluorescence staining

In vitro cultured cells were fixed with 4% paraformaldehyde for 10 minutes and permeabilized with 0.2% Triton X-100 for 5 minutes. After blocking with 2% BSA in PBS containing 0.5% Tween-20, cells were stained with RTF1 (Sigma) primary antibody, followed by blotting with fluorescent conjugated secondary antibody. Nuclei were labeled with DAPI (Sigma). Fluorescence images were acquired using a motorized Nikon Ti inverted microscope equipped with a Yokogawa CSU-W1 spinning disk confocal head with a 50 μm pinhole size, an Andor Zyla 4.2 plus sCMOS camera, Toptica iChrome MLE 4-color multi-laser engine (488, 515, 561 and 640 nm), SOLA 395 engine widefield illuminator, and Nikon motorized stage with a PI 250 mm piezo insert, using a Plan Apo 100x/1.45 DIC objective. Fluorescence imaging was performed using a Chroma quad multipass dichroic mirror and Chroma 525/36 nm bandpass emission filter. The acquisition software was NIS Elements AR 5.02.

### Retroviral production and transduction

Mouse RTF1 WT and an HMD mutant with 8 alanine replacements at conserved amino acids (D194, D197, E206, E208, E210, E214, E216, and R225) were cloned into retroviral vector MIGR-IRES-GFP with a flag tag at the N-terminal. Mouse RORγt fused with BFP at the C-terminal was cloned into MIGR. Vectors were co-transfected into 293T cells along with 10A1 packaging plasmid for retrovirus production. The resulting viral supernatants were collected and directly used for transduction. To transduce CD4^+^ T cells, naïve T cells were cultured under Th0 conditions for 24 hours. Cells were then centrifuged with viral supernatant containing 1 mg/mL polybrene at 2500 rpm for 2 hours at 37 °C. Subsequently, the media was replaced with Th17 polarization conditions, and cells were cultured for an additional 3 days. Virus-transduced cells were gated as GFP^+^ or BFP^+^ populations and analyzed by flow cytometry.

### Gut lamina propria preparation

Gut tissues were dissected and treated with 1 mM DTT at room temperature for 10 minutes, and 5 mM EDTA at 37 °C for 20 minutes to remove epithelial cells. The dissociation of tissues was performed using a digestion buffer (RPMI, 1 mg/mL Liberase, 100 μg/mL DNase I, 5% FBS) with continuous stirring at 37 °C for 30 minutes. Mononuclear cells were isolated from the interface of a 30%/70% Percoll gradient (GE Healthcare). The collected cells were then analyzed by flow cytometry.

### Adoptive transfer and tamoxifen treatment

CD4^+^CD25^−^CD62L^hi^CD44^low^ naïve T cells were sort purified from WT B6.CD45.1, *Rtf1^UBC^*.CD45.2, or RNF40^Rosa^.CD45.2 mice using FACS sort, and then mixed at a 1:1 ratio. Subsequently, 0.5 million cells were adoptively transferred into each Rag1-KO recipient mouse through intraperitoneal (i.p.) injection. The recipient mice received tamoxifen treatment at a dosage of 75 mg/kg through i.p. administration once a day for a total of 5 consecutive days. After two weeks, spleen and ileum tissues were collected, and lamina propria lymphocytes were dissociated and analyzed by flow cytometry.

### RNA sequencing and analysis

Naïve CD4+ T cells were purified from *Rtf1^UBC^* mice and cultured under Th17 conditions for 3 days in the presence of either DMSO or 4-hydroxytamoxifen. Total RNA extraction and purification were performed using the RNeasy mini kit (Qiagen) following the manufacturer’s instructions. The purified RNA samples were then sent to Harvard Biopolymers Core for library preparation and deep sequencing. The sequencing reads were aligned to the Mus musculus genome (version mm10) using STAR alignment software tools. Calculation of read counts was carried out using featureCounts. Gene expression normalization was performed using DESeq2, and differential expression genes were depicted in a scatter plot. The RNAseq and Array data were analyzed separately due to inherent differences between the two platforms. RNAseq DEGs were identified with p-value(padj) <0.05, log2FC >1 or log2FC < -1. Array DEGs were directly downloaded from Enrichr website (https://maayanlab.cloud/Enrichr/#meta!meta=GSE27241). The comparison was made at the level of DEGs identified in each dataset independently. The Venn diagram was generated by the R package VennDiagram. The heatmap clustering was generated using the R package pheatmap.

### CUT&RUN

CUT&RUN of in vitro-differentiated Th17 cells was performed using Epicypher’s CUTANA™ ChIC/CUT&RUN Kit (#141-048) as per the manufacturer’s recommendations with slight modifications. Briefly, nuclei were prepared using a nuclear extraction buffer (20 mM HEPES pH 7.9, 10 mM KCl, 0.1% Triton X-100, 20% Glycerol, 0.5 mM Spermidine, 1x Roche cOmplete™, Mini, EDTA-free Protease Inhibitor) from in vitro-differentiated Th17 cells. 500k nuclei were used as the starting input amount for each reaction with three replicates for each RTF1 (Bethyl Laboratories #A300-179A, 0.5µg per reaction) and H2Bub1 (Cell Signaling Technology #5546, 0.5µg per reaction) performed. Additional reactions were performed as recommended using 1µL of H3K4me3 as a positive control (EpiCypher #13-0041) and IgG as a negative control (EpiCypher #13-0042). Antibody binding for all samples was performed overnight at 4 °C on a nutator after which 2.5 µL/reactions of pAG-MNase and 1 µL/reaction of 100 mM calcium chloride were added to cleave and release antibody-bound DNA. Library preparation was performed using NEBNext® Ultra™ II DNA Library Prep Kit for Illumina® (NEB #E7645S) as per the manufacturer’s recommendations and indexed using NEBNext® Multiplex Oligos for Illumina® (96 Unique Dual Index Primer Pairs) (NEB #E6440S). Final libraries were analyzed using Agilent’s Bioanalyzer system (Agilent High Sensitivity DNA Kit Cat #50674626) before being sequenced using Illumina’s NextSeq2000. Primary analysis was performed using nf-core/cutandrun (Cheshire et al., 2024) with alignment to the mm10 reference genome and SEACR (stringent) for peak calling (Meers et al., 2019). Downstream analysis was performed in R using peaks assigned in all three for each RTF1 and H2Bub1 and overlapping with EPDnew promoters (Dreos et al., 2013).

### Statistical analysis

The data is presented as mean ± SD, and unless otherwise specified, all presented data represent results from a minimum of three independent replicates. Statistical analysis was conducted using Prism 8 (GraphPad), and a two-tailed Student’s t-test was applied for statistical comparisons. Significance levels are denoted as follows: \**P* < 0.05, \*\**P* < 0.01, \*\*\**P* < 0.001, \*\*\*\**P* < 0.0001.

### Data availability

The RNA-sequencing data have been deposited in the Gene Expression Omnibus under accession code GSE269436. CUT&RUN data have been deposited and publicly available at GEO accession number GSE269642. All other data supporting the findings of this study are available from the corresponding author upon reasonable request.

## Results

### A screen for modulators of RORγt identifies RTF1

With *Drosophila* S2 cell lines, we previously carried out two unbiased screens, a small molecule screen and a double-stranded RNA (dsRNAs) screen, to identify chemical and genetic modulators of RORγt activity (Huh et al., 2011). The former led to the identification of the first small molecule inhibitor targeting RORγt with high specificity. We then performed both primary and secondary genetic screens with S2 cell reporters monitoring the transcription activities of related nuclear hormone receptors (NHR), including Daf12, an NHR that senses sterol derivative levels in *C. elegans* (Antebi et al., 2000) as the control. From this screen, *Drosophila Rtf1* was identified as a positive regulator of RORγt (**Fig. S1A-B**).

### *RTF1* genetic variants are associated with the risk of ulcerative colitis

*RTF1* has previously been reported to be associated with ulcerative colitis risk in multiple genome-wide association studies (GWAS) (Jostins et al., 2012; Lange et al., 2017; Liu et al., 2015, 2023). The first variant, rs28374715, is located in the upstream region of the *RTF1* gene, in an intronic region of the *CHP1* gene, and is associated with an increased risk of ulcerative colitis (*p* = 2.43E-08, OR = 1.08 (1.04-1.13) in European ancestry individuals) (Jostins et al., 2012) (**Fig. S2**). The second variant, rs2777491, is located within the second intron of *RTF1*, and unlike the first one, is associated with a decreased risk of ulcerative colitis (*p* = 8.32E-08, OR = 0.94 [0.91-0.96]) in European ancestry individuals (Liu et al., 2023) (**Fig. S2**). We next investigated whether these two GWAS variants are also associated with *RTF1* expression quantitative trait loci (eQTLs). Intriguingly, we found *RTF1* eQTLs colocalizing near the second variant, rs2777491 (H4 = 99.6%) in whole-blood cells (**Fig. S2**). These genetic analysis data indicate an interesting possibility that altered *RTF1* transcription may underlie its associated genetic risk of developing ulcerative colitis. RTF1 was identified as a regulator of RORγt (**Fig. S1A-B**). Given that RORγt serves as the master regulator of Th17 cells differentiation and function, and Th17 cells have been extensively implicated in the pathogenesis of inflammatory bowel diseases (IBD), we sought to investigate whether RTF1 plays a critical role in regulating Th17 cells.

### RTF1 regulates T cell development

We first assessed whether RTF1 proteins are expressed in T cells in mice. Western blot analyses identified its expression across different developmental stages, encompassing naïve, effector T cells, and thymic-derived regulatory T cells (nTreg). Furthermore, RTF1 protein is robustly expressed in in vitro cultured T cells, including Th1, Th2, Th17, and peripherally induced Treg cells (iTreg) (**Fig. S3A**). Next, we generated a conditional knockout mouse line, *Rtf1^CD4^*, by crossing mice with a loxP-flanked RTF1 allele (Austin et al., 2004) with those expressing CD4-Cre. Total CD4^+^ T cells isolated from *Rtf1^CD4^* mice showed a significant reduction of RTF1 protein (**Fig. S3B**). In the thymus, *Rtf1^CD4^*mice were found to have similar frequencies of CD4^+^ and CD8^+^ T cell populations to those of wild-type (WT) mice. However, we observed a substantially increased percentage of an immature double-negative T cell population in the spleens (**Fig. S3C**), accompanied by a significant reduction in total thymic and splenic T cell numbers (**Fig. S3D**). Subsequently, the loss of RTF1 led to a significant decrease in both naïve CD4^+^ and CD8^+^ T cell percentages in spleens (**Fig. S3E-G**). These results suggest that RTF1 plays a crucial role in regulating thymic T cell development by likely supporting T cells’ survival or proliferation; without it, overall T cell numbers in the periphery are decreased.

### RTF1 regulates Th17 differentiation in vitro

To circumvent the developmental impact of RTF1 deletion in CD4^+^ T cells and probe its role in mature CD4^+^ T cell subsets, we generated an inducible knockout mouse line, *Rtf1^UBC^*, by breeding *Rtf1*-floxed mice with transgenic mice expressing ubiquitously active tamoxifen-inducible Cre (UBC-Cre-ERT2). By treating cells isolated from the *Rtf1^UBC^* mouse line with the synthetic estrogen receptor ligand, 4-hydroxytamoxifen (4OHT), we successfully removed the *Rtf1* gene. Indeed, in vitro cultured naïve CD4^+^ T cells isolated from *Rtf1^UBC^*mice displayed a marked reduction in RTF1 protein levels following 4OHT treatment (**Fig. 1A**). RTF1 protein is predominantly localized within the nucleus of control T cells, but its level is substantially reduced in 4OHT-treated cells (**Fig. 1B**). While in vitro-differentiated T helper cell subsets express a high level of RTF1 (**Fig. S3A**), RTF1 removal preferentially affects T cells cultured under Th17 polarizing conditions with a pronounced reduction in interleukin-17a (IL-17a) expression (**Fig. 1C**). In contrast, RTF1 deletion did not impact Th1 or iTreg differentiation, assessed by cytokine IFNγ, and transcription factor forkhead box P3 (FOXP3) expression, respectively (**Fig. 1C**). We also isolated naïve CD4^+^ T cells from *Rtf1-*floxed mice and transduced them with retroviruses expressing GFP or GFP-Cre, followed by culturing them under a Th17-polarizing condition. Consistent with *Rtf1^UBC^* cell data, Cre-transduced cells exhibited a substantial reduction in IL-17a expression compared to those transduced with the GFP control virus (**Fig. S4A**). Therefore, RTF1 deletion in naïve CD4^+^ T cells selectively impairs Th17 cells by affecting their survival or proliferation without affecting iTreg differentiation.

**Figure 1.**
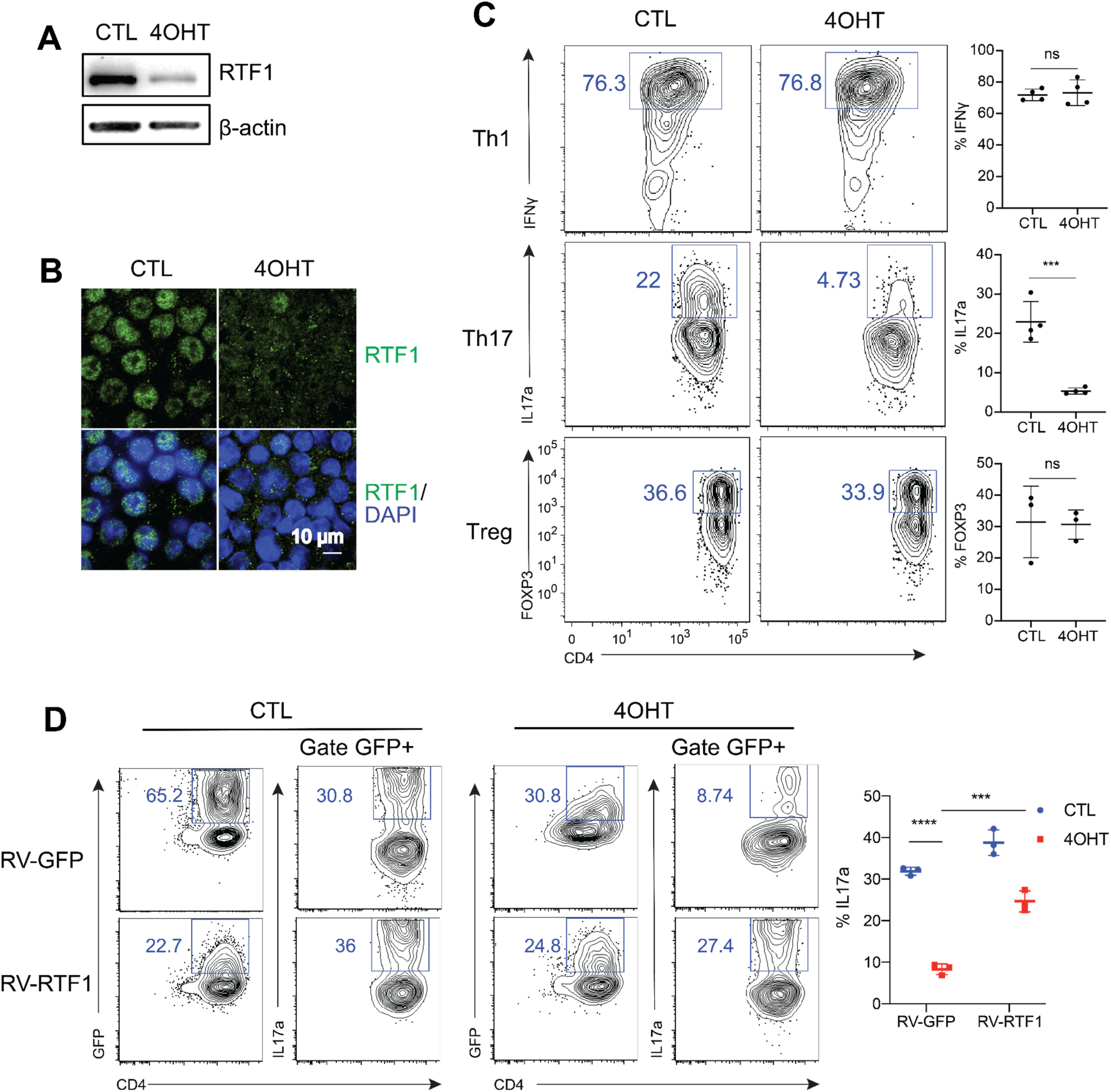
RTF1 is required for Th17 cell differentiation in vitro. Naïve CD4^+^ T cells from *Ubc^Cre-ERT2^Rtf1^fl/fl^ (Rtf1^UBC^)* mice were isolated, and in vitro cultures were treated with the estrogen receptor agonist 4-hydroxytamoxifen (4OHT) or DMSO as a control (CTL). **A** and **B**. Immunoblotting and Immunostaining of RTF1 in cells cultured under Th0 conditions (anti-CD3, anti-CD28, IL-2). β-actin served as a loading control. **C**. Flow cytometric and statistical analysis of cells cultured under Th1 (anti-CD3, anti-CD28, IL-2, IL-12), Th17 (anti-CD3, anti-CD28, IL-6, TGFβ) or iTreg (anti-CD3, anti-CD28, IL-2, TGFβ) polarizing conditions for three days, followed by stimulation with PMA, ionomycin, and Golgi plug for 4 hours to assess cytokine expression through intracellular staining. **D**. Flow cytometric and statistical analysis of cells cultured under Th17 conditions and transduced with retrovirus carrying GFP (RV-GFP) or RTF1 (RV-RTF1). Experiments were conducted using 5 to 8-week-old mice. The data depict analysis from 3 to 4 mice per genotype (mean ± SD; unpaired *t*-test). ns = not significant. \*\*\**P* < 0.001, \*\*\*\**P* < 0.0001.

We then tested whether RTF1 overexpression restores IL-17a production from T cells lacking the *Rtf1* gene. Naïve CD4^+^ T cells isolated from *Rtf1^UBC^* mice were transduced with control (RV-GFP) or RTF1 expressing (RV-RTF1) retroviruses. Even after 4OHT treatment, RV-RTF1-T cells were found with significantly enhanced IL-17a expression, compared to RV-GFP-T cells (**Fig. 1D**). Consistent with its critical role as a master transcription factor for Th17 cell differentiation (Ivanov et al., 2006), ectopic expression of RORγt in T cells (RV-RORγt) produces IL-17a even though they were cultured under a Th0 non-polarizing condition (**Fig. S4B**). Importantly, RTF1 deficiency (4OHT-treated cells) significantly compromised RV-RORγt’s ability to enhance IL-17a expression (**Fig. S4B**). Global transcriptional profiling analyses comparing RTF1-expressing and -depleted cells revealed that RTF1 promotes the expression of both IL-17a and IL-17f, Th17 cell signature genes, without affecting *Rorc* expression. These data suggest that RTF1 may directly regulate the expression of RORγt target genes (**Fig. S5A**). Further supporting this notion, differentially expressed genes (DEGs) between control and RTF1-deficient cells substantially overlapped with the DEGs between control and *Rorc* knockout cells (**Fig. S5B**). Furthermore, a majority of those genes commonly regulated by both RORγt and RTF1 display a similar expression pattern in cultured Th17 cells (**Fig. S5C, Supplemental Table 1**). Thus, RTF1, like RORγt, supports a key transcription program of Th17 cells in vitro, but likely functions downstream of RORγt.

### Loss of RTF1 impairs Th17 development in vivo

To understand the role of RTF1 in CD4^+^ T cells in vivo, we isolated naïve CD4^+^ T cells from WT (CD45.1) mice and RTF1^UBC^ mice (CD45.2). Isolated cells were then mixed at a 1:1 ratio and adoptively transferred into Rag1-knockout recipient mice (**Fig. 2A-B**), followed by tamoxifen treatment to induce RTF1 deletion in RTF1^UBC^ cells. Two weeks post-transfer, a notable reduction in the number of RTF1^UBC^ cells, compared to WT cells, was evident in both spleens and ilea of the recipient mice, indicating that RTF1-depleted cells are outcompeted by RTF1-non-depleted cells (**Fig. 2C**). Consistent with in vitro cell culture data (**Fig. 1C**), a significantly lower proportion of RTF1^UBC^ cells underwent differentiation into Th17 cells compared to WT cells (**Fig. 2D**). To specifically assess the impact of RTF1 deletion in differentiated Th17 cells, we next generated RTF1 deficiency in IL-17a-expressing cells by crossing IL17a-Cre (RTF1^IL17a^) and RTF1-floxed mice. Under a steady-state condition, the ilea of RTF1^IL17a^ mice exhibited significantly fewer Th17 cells than control mice (**Fig. 2E-F**). These data confirm the key function of RTF1 in in vitro Th17 cell generation and maintenance.

**Figure 2.**
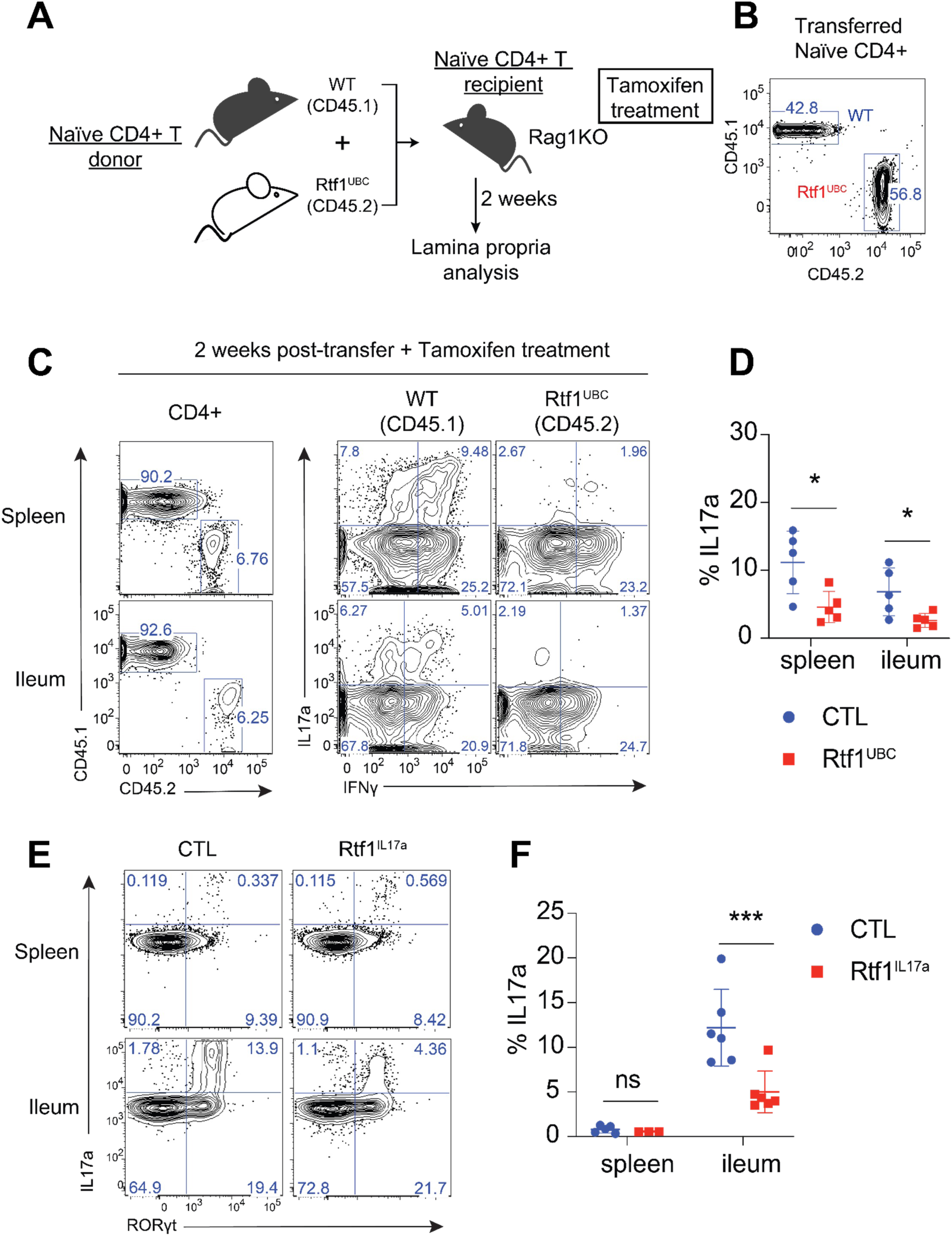
RTF1 controls Th17 cell development in vivo. Naïve CD4^+^ T cells from WT CD45.1 and *Ubc^Cre-ERT2^Rtf1^fl/fl^ (Rtf1^UBC^,* CD45.2) mice were mixed at a 1:1 ratio and adoptively transferred into *Rag1* knockout recipients (*Rag1KO*). The recipients received 75 mg/kg tamoxifen via intraperitoneal injection for 5 consecutive days. Two weeks later, the splenocytes and ileal lamina propria T cells were isolated and analyzed by flow cytometry. **A**. Overview of the experimental design. **B**. Pre-transfer status of WT and *Rtf1^UBC^* T cells. **C**. Representative flow cytometric plots displaying transferred WT and *Rtf1^UB^*^C^ T cells at day 14 post-inoculation. **D**. Statistical analysis of Th17 cells from spleen and ileum. **E** and **F**. Flow cytometric and statistical analysis of Th17 cells from spleen and ileum of IL-17A^Cre^ control (CTL) or IL-17A^Cre^RTF1^fl/fl^ (RTF1^IL17a^) mice at steady state. In vivo experiments were performed once with a total of 5 or 6 mice per group. The results shown are representative of this single experiment, which was consistent across different iterations. Data represent 5 to 6 mice per genotype analyzed. Mice aged 5 to 8 weeks were used. Statistical data are the mean ± SD; unpaired *t*-test. ns = not significant. \**P* < 0.05, \*\*\**P* < 0.001.

### RTF1 regulates the H2B mono-ubiquitination in Th17 cells

In yeast, RTF1 enhances H2Bub1, and its histone modification domain (HMD) directly interacts with the ubiquitin-conjugating enzyme RAD6 (Oss et al., 2016), a homolog of mammalian RNF40. Consistent with these data, deleting RTF1 reduced H2Bub1 levels in Th17 cells (**Fig. 3A**). Compared to control virus-transduced cells with or without 4OHT treatment, cells transduced with RV-RTF1 (WT) were found with restored H2Bub1 expression. On the other hand, overexpression of RTF1 mutant (mHMD) carrying eight conserved amino acid substitutions with alanine within the HMD domain failed to achieve the same effect in the RTF1-deficient cells (**Fig. 3B-C**). Of note, both WT and mHMD RTF1 levels are expressed at comparable levels (**Fig. 3D**). Furthermore, *Rtf1*-mHMD expression, unlike its WT counterpart, no longer restored IL-17a expression in Th17 cells devoid of RTF1 (**Fig. 3E**), highlighting the requirement of the functional HMD domain and associated H2Bub1-promoting activity for IL-17a expression.

**Figure 3.**
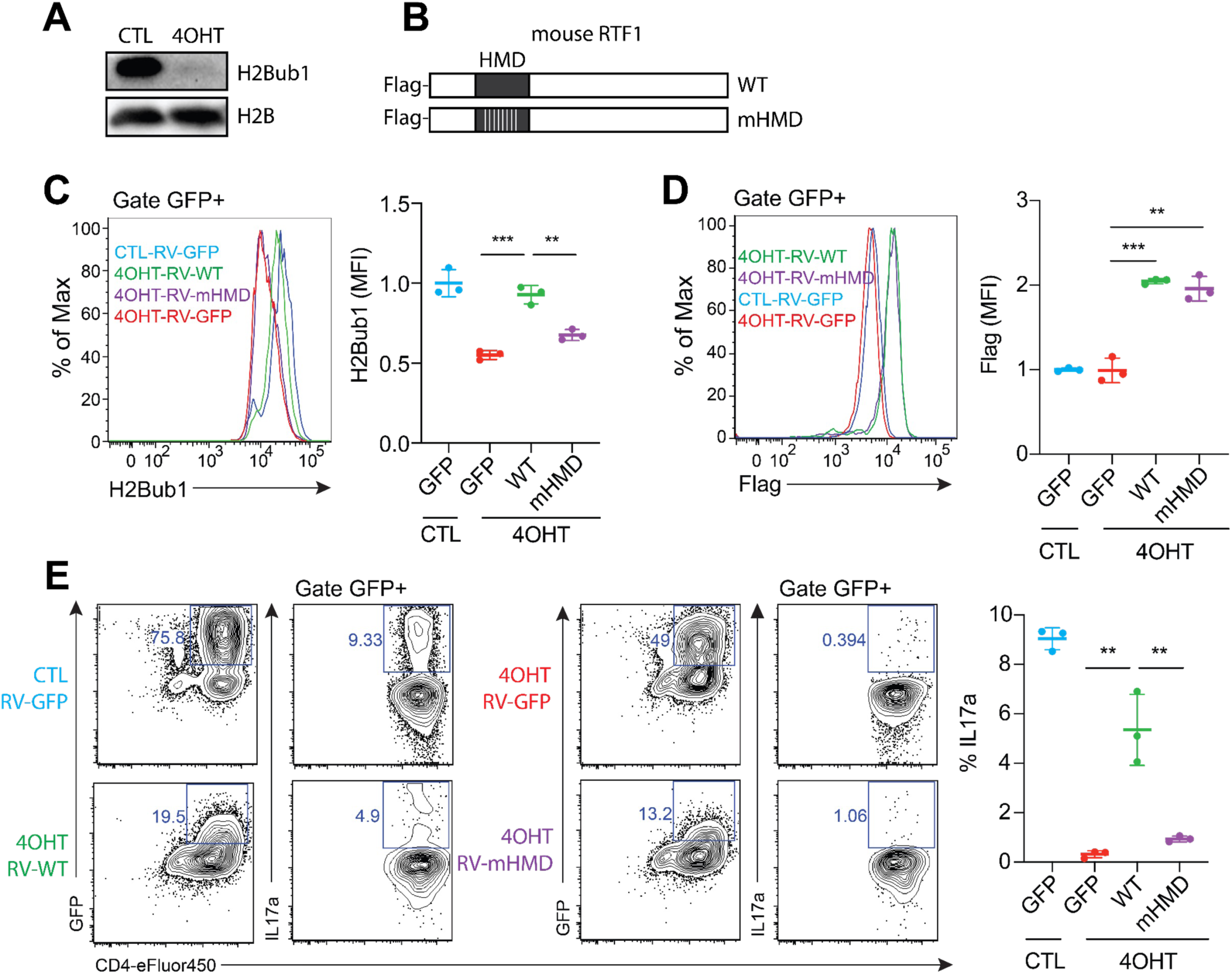
RTF1-dependent H2B monoubiquitination regulates Th17 differentiation. **A**. Immunoblotting analysis of H2B monoubiquitination (H2Bub1) and total H2B in cultured Th17 cells isolated from *Rtf1^UB^*^C^ mice treated with DMSO (CTL) or 4-hydroxytamoxifen (4OHT). **B**. Schematic representation of wild-type *Rtf1* (WT) and the mutant form of *Rtf1* (mHMD) with 8 conserved amino acids in the HMD domain replaced by alanine. **C**-**F**. Retroviral transduction of cultured Th17 cells isolated from *Rtf1^UBC^*mice. Cells were transduced with retroviruses carrying GFP (RV-GFP), WT (RV-WT), or mHMD (RV-mHMD) constructs and analyzed 72 hours post-transduction. **C**. Flow cytometric and statistical analysis of H2Bub1 in retroviral-transduced Th17 cells cultured with DMSO (CTL) or 4-hydroxytamoxifen (4OHT). **D**. Flow cytometric and statistical analysis of the flag tag for monitoring the expression levels of retrovirally transduced constructs in Th17 cells cultured with DMSO (CTL) or 4-hydroxytamoxifen (4OHT). The control GFP construct does not carry a flag tag, while both RTF1 and mHMD constructs carry a flag tag at the N-terminus. **E**. Flow cytometric and statistical analysis of IL-17a expression in retrovirally transduced Th17 cells cultured with DMSO (CTL) or 4-hydroxytamoxifen (4OHT). Data represent three independently replicated samples (mean ± SD; unpaired *t*-test). \*\**P* < 0.05, \*\*\**P* < 0.001.

### H2Bub1 is required for Th17 cell differentiation

To directly assess if H2Bub1 supports Th17 cell differentiation, we generated an *Rnf40*-floxed mouse line and established a Rosa-Cre-ERT2-driven, inducible RNF40 (*Rnf40^Rosa^*) mouse model. RNF40 is a key component of the ubiquitin ligase complex (RNF20/RNF40), responsible for H2Bub1 at lysine 120 (Cole et al., 2015). Like RTF1, RNF40 deficiency resulted in impaired H2Bub1 (**Fig. 4A**) and, importantly, compromised Th17 cell differentiation in vitro without affecting iTreg differentiation (**Fig. 4B**). However, we found that RNF40 deletion also led to reduced Th1 differentiation, indicating H2Bub1 may play a role in supporting both Th1 and Th17 programs, while RTF1 has a more specific impact on Th17 cells. Using the same adoptive transfer model as depicted in Figure 2A, we next tested whether RNF40 deletion phenocopies RTF1 removal in T cells in vivo. Supporting this hypothesis, compared to WT control, we observed reduced cell numbers and impaired Th17 cell levels in RNF40-deficient T cells in recipient mice (**Fig. 4C-E**), further underscoring the critical function of RTF1 and RNF40-mediated H2Bub1 in Th17 cells.

**Figure 4.**
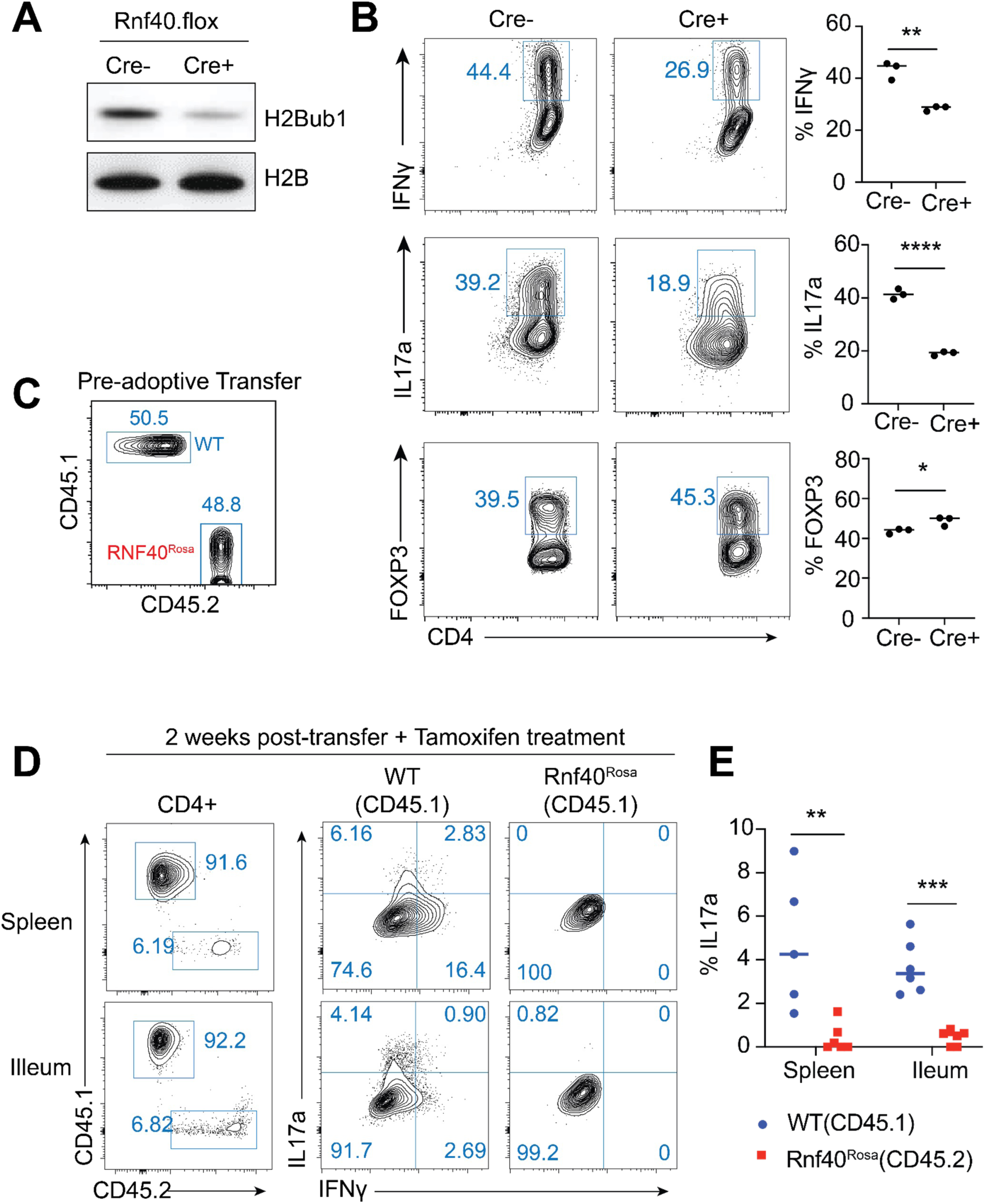
H2Bub1 ubiquitin ligase RNF40 is essential for Th17 differentiation in vitro and in vivo. Naïve CD4^+^ T cells from *Rnf40^fl/f^*^l^ (Cre-) or ROSA^Cre-ERT2^*Rnf40^fl/fl^* (Cre+) mice, and in vitro cultures were treated with 4-hydroxytamoxifen (4OHT). **A**. Immunoblotting analysis of H2Bub1 and total H2B in cultured Th17 cells. **B.** Flow cytometric and statistical analysis of cells cultured under Th1 (anti-CD3, anti-CD28, IL-2, IL-12), Th17 (anti-CD3, anti-CD28, IL-6, TGFβ) or iTreg (anti-CD3, anti-CD28, IL-2, TGFβ) polarization conditions for three days, followed by stimulation with PMA, ionomycin, and Golgi plug for 4 hours to assess cytokine expression through intracellular staining. **C-E.** Adoptive transfer experiments were performed the same way as in Figure 2A-D, with a 1:1 mixture of cells isolated from WT CD45.1 and *RNF40^Rosa^*.CD45.2 mice. **C**. Pre-transfer status of WT and *RNF40^Rosa^* T cells. **D**. Representative flow cytometric plots displaying transferred WT and *RNF40^Rosa^* T cells at day 14 post-inoculation. **E**. Statistical analysis of Th17 cells from spleen and ileum. In vivo experiments were performed once with a total of 5 mice per group. The results shown are representative of this single experiment, which was consistent across different iterations. Mice aged 5 to 8 weeks were used. Statistical data are the mean ± SD; unpaired *t*-test. \**P* < 0.05, \*\**P* < 0.01, \*\*\**P* < 0.001, \*\*\*\**P* < 0.0001.

### RTF1 and H2Bub1 colocalize on known Th17 regulators

We next performed CUT&RUN with in vitro-derived Th17 cells to examine whether RTF1 and H2Bub1 bind to similar loci in Th17 cell-associated genes (**Fig. S6A**). Binding peaks for both RTF1 and H2Bub1 were strongly enriched at gene promoter regions and more than 70% of both factors’ called peaks were assigned within 1kb of gene promoters with detectable enrichment of signal distribution around transcriptional start site (TSS) regions (± 3kb) (**Fig. 5A, B**).

**Figure 5.**
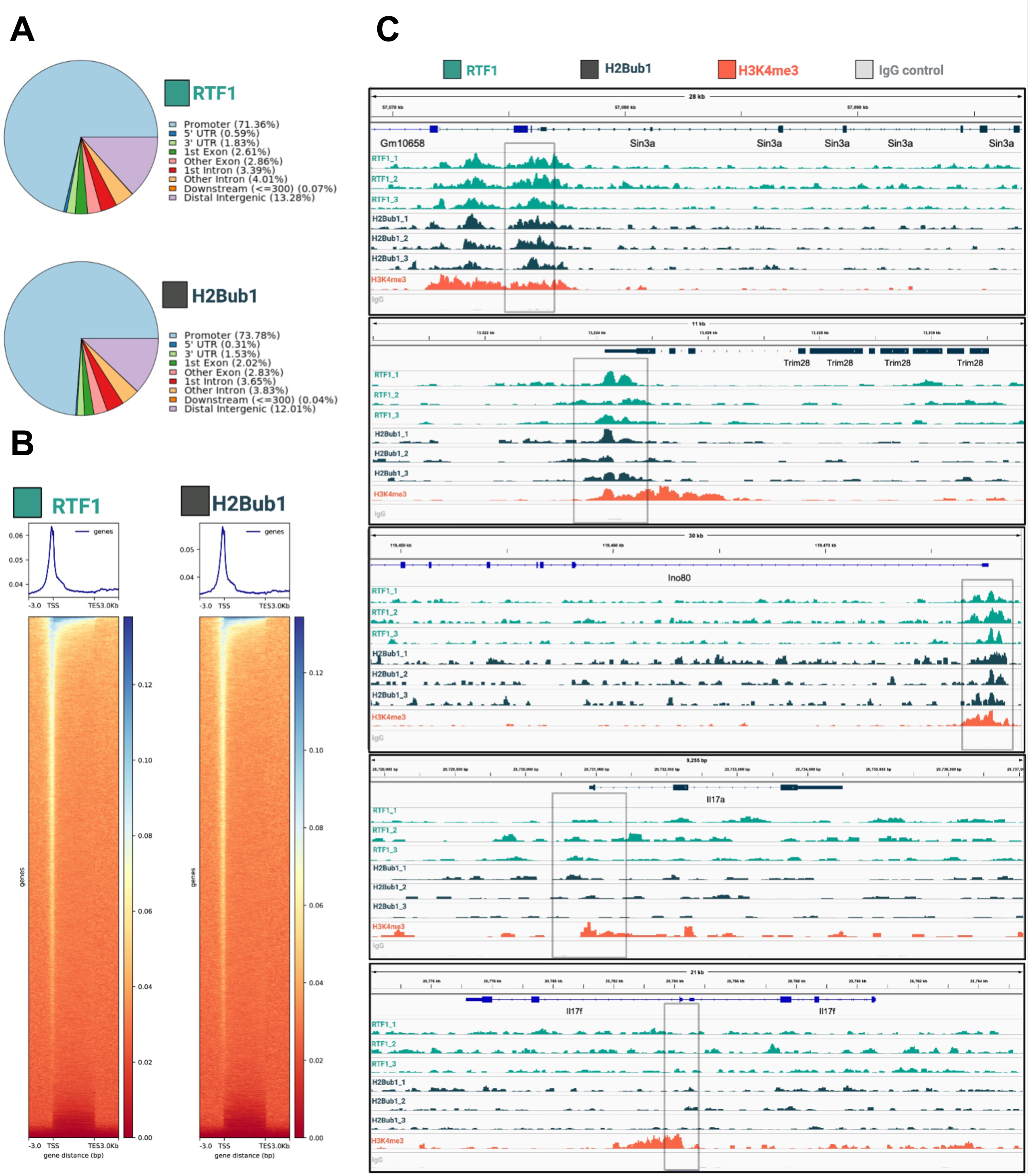
RTF1 and H2Bub1 colocalize on known Th17 cell regulators. **A**. Distribution of genomic features for RTF1 and H2Bub1 with more than 70% of peaks overlapping with gene promoter regions. **B**. Enrichment of signal distribution of RTF1 and H2Bub1 +/-3kb relative to TSS. Y axis denotes peak count frequency. **C**. IGV panels of all three replicates of RTF1, H2Bub1, and control H3K4me3/IgG binding are shown for *Sin3a, Trim28, and Ino80*. Notable absence of RTF1 and H2Bub1 binding for *Il17a* and *Il17f.* Annotated gene promoters are boxed in gray.

Consistent with RTF1’s major function in enhancing H2B mono-ubiquitylation, we found 75% of all H2Bub1 peaks overlapped with RTF1 binding peaks (**Fig. S6B, Supplemental Table 2**). We then performed gene ontology analyses with those genes of which promoter regions were occupied by both RTF1 and H2Bub1 and found the most highly enriched gene set term was histone modification (**Fig. S6C**). This subset includes genes regulating nearly all aspects of chromatin modification and genomic integrity critical for cell lineage commitment and maintenance such as *Ino80, Ezh2, Jarid2, Kdm6b/Jmjd3, Satb1, Trim28,* and *Sin3a* (**Fig. 5C**). Specifically, *Sin3a,* a known regulator of Stat3 transcription (Icardi et al., 2012), has been shown to balance *Il17a* and *Foxp3* expression in CD4^+^ T cells (Perucho et al., 2023). Cells deficient in Sin3A fail to properly upregulate *Il17a* (as well as *Il17f*, *Il23r*, and *Il22*), but instead exhibit an IL2^+^ and FOXP3^+^ phenotype (Perucho et al., 2023). Similarly, TRIM28 (KAP1/TIF1β) was shown to function as a positive regulator of Th17 differentiation with loss of *Trim28* leading to a reduction in *Il17a* and *Il17f* expression levels while increasing *Foxp3* expression (Jiang et al., 2018). Reminiscent of RTF1 phenotypes, loss of *Trim28* did not impact *Rorc* expression. Mechanistically, it was reported that STAT3 first recruits TRIM28 to key loci, which allows for RORγt recruitment to support IL-17 production (Jiang et al., 2018).

We also found that RTF1 co-localized with H2Bub1 at promoter regions of classical regulators for Th17 differentiation pathways, such as *Stat3, Batf, cMaf, Junb, Hif1a,* and *Rora*, among others (**Fig. S6D**). Notably, we did not detect RTF1 or H2Bub1 binding to the *Rorc* promoter region despite a strong H3K4me3 signal (**Fig. S6D**). This finding is consistent with our gene expression data in which *Rorc* expression was not impacted in RTF1-deficient cells (**Fig. S5A**). RTF1 and H2Bub1 binding was also absent in *Il17a* and *Il17f* promoters (**Fig. 5C**), suggesting the enhanced expression of IL-17a and IL-17f by RTF1 and H2Bub1 does not require direct binding to their promoter regions. Taken together, RTF1 and H2Bub1 function as key epigenetic regulators in Th17 cell differentiation and function by likely modulating chromatin recruitment, remodeling, and accessibility necessary for the Th17 cell program.

## Discussion

Th17 cells, a subset of CD4+ T cells, are known for their role in promoting inflammatory responses through the production of effector cytokines, particularly IL-17a and IL-17f. GWAS findings have linked genes involved in Th17 cell differentiation and function, such as IL-23R and STAT3, with altered IBD susceptibility (Momozawa et al., 2011). Numerous studies have demonstrated an increased presence of Th17 cells and elevated IL-17 levels in the intestinal mucosa of IBD patients (Fujino et al., 2003; Jiang et al., 2014; Sugihara et al., 2010; Lloyd-Price et al., 2019). Moreover, animal models and cell culture experiments have provided further insights into the inflammatory roles of Th17 cells in driving intestinal inflammation (Feng et al., 2011; Ivanov et al., 2006; Kullberg et al., 2006; Lu et al., 2015; Ishigame et al., 2009), suggesting an intricate connection between genetic variants, the Th17 cell pathway, and IBD pathogenesis.

In this study, we provided a series of in vitro and in vivo evidence supporting the critical function of RTF1 in Th17 cell differentiation and function. Our eQTL analyses, along with associated genetic risks of RTF1 variants with ulcerative colitis, proposed a plausible mechanism by which alteration of RTF1 expression may contribute to UC and other IBD phenotypes through modulating the Th17 cell program. Intriguingly, another genetic RTF1 variant, rs12440045, exhibits a robust association with neutrophil counts (Astle et al., 2016). Given that IL-17a plays a key role in neutrophil mobilization and activation (Zenobia and Hajishengallis, 2014), and considering that neutrophil infiltration is a hallmark of UC and disease severity (Fournier and Parkos, 2012; Friedrich et al., 2021; Wéra et al., 2016), RTF1 may also orchestrate the cross-talk between Th17 cells and neutrophils in the context of IBD.

RTF1 is also a known constituent of the PAF1C complex in yeast, along with Paf1, Ctr9, and Leo1, where this assembly governs various steps of RNA polymerase II-dependent transcription, encompassing transcriptional initiation, elongation, and RNA processing (Jaehning, 2010). Both human and mouse RTF1, however, are also known to operate independently of the PAF1C (Cao et al., 2015; Parnas et al., 2015). For instance, RTF1 and PAF1C regulate the expression of distinct gene sets in cultured human cancer cell lines (Cao et al., 2015). In mouse macrophages, RTF1 doesn’t physically interact with PAF1C (Parnas et al., 2015). Furthermore, mouse CTR9 acts as a suppressor of Th17 differentiation (Yoo et al., 2014). These data suggest that RTF1’s function in Th17 cells differs from its possible role, if any, in PAF1C.

Two key domains of the RTF1 protein, the HMD and Plus3 domains are evolutionarily conserved from yeast to humans (Jong et al., 2008; Warner et al., 2007). Indeed, the RTF1 HMD (a critical domain for histone modification) from various species are capable of compensating for the defects resulting from HMD deletion in yeast (Piro et al., 2012). Conversely, the Plus3 domain has been demonstrated to bind single-stranded DNA and the phosphorylated form of Spt5 (Jong et al., 2008; Wier et al., 2013). In murine CD4^+^ T cells, our data likely support the idea that RTF1 possesses dual roles in promoting both cell proliferation and Th17 differentiation.

H2B is co-transcriptionally ubiquitylated, and its levels correlate with RNA polymerase II elongation rates (Fuchs and Oren, 2013; Kim et al., 2010). H2Bub1 is enriched at active transcription sites (Minsky et al., 2008), and it is essential for di-and tri-methylation of H3K4 and H3K79, marking the active chromatin regions and, thus, contributing to proper silencing and the establishment of heterochromatin regions (Krogan et al., 2002; Nguyen and Zhang, 2011; Smolle and Workman, 2013). Van Oss et al. demonstrated that RTF1 directly interacts with the ubiquitin conjugate RAD6, promoting H2Bub1 and enhancing nucleosome stability in yeast (Oss et al., 2016). In line with this, our mouse CD4^+^ T cell data indicated that RTF1 deletion substantially diminished H2Bub1 in Th17 cells without affecting the overall H2B level. H2Bub1 levels and the corresponding IL-17a expression in Th17 cells could be restored through RTF1 overexpression, underscoring the positive regulatory role of RTF1-dependent H2Bub1 in Th17 cell development.

The transcriptional regulatory program governing Th17 cell differentiation and effector function is primarily directed by the nuclear transcription factor RORγt (Ivanov et al., 2006; Manel et al., 2008). Additionally, other transcription factors, such as HIF1a, STAT3, and c-MAF, play key roles (Dang et al., 2011; Poholek et al., 2020; Ciofani et al., 2012) supporting the Th17 cell program. We found that RTF1 is not directly localized at the *Il-17* gene locus (**Fig. 5C**). In addition, RTF1 does not directly control *Rorc* expression (**Fig. S5**), even though it is crucial for RORγt activity (**Fig. S1**) and RORγt-driven Th17 differentiation (**Fig. S4B**). In addition to supporting RORγt activity as a transcription factor, RTF1, and H2Bub1, may potentiate the Th17 cell program through the action of other transcription factors, such as HIF1a and STAT3.

Consistently, their promoter regions are bound by RTF1 and H2Bub1. Interestingly, *Sin3a*, a necessary component for the hypoxia response (Tiana et al., 2018), is also highly associated with both RTF1 and H2Bub1 binding sites. These data indicate the presence of multiple mechanisms by which RTF1 promotes Th17 cell function. Of note, RORγt also plays a key modulatory function in a subset of innate immune cells, such as ILC3s and MAIT cells and innate-like γδT cells. Therefore, it will be crucial to examine whether RTF1 also supports the function and differentiation of these RORγt-expressing innate immune cells.

In summary, our study identified that RTF1 enhances Th17 differentiation and function, likely with the help of other Th17 cell-associated transcription factors, but acting downstream or independent of RORγt. We found RTF1 along with RNF40 may serve as an epigenetic regulator enhancing H2Bub1 levels and boosting the Th17 program. Our study supports the notion, therefore, that epigenetic interventions, such as targeting H2Bub1, will likely serve as viable therapeutic strategies to modulate Th17 cell function for addressing various autoimmune and inflammatory diseases.

## Acknowledgments

We thank Lucinda Tam and Fermin Gallardo-Chang for helping with the design and maintenance of *Rnf40* knockout mice. This study was supported by the NIH grants (R01 DK106351 and R01 DK110559-08) to J.R.H. The authors have no conflicting financial interests.

---

**Figure S1.**
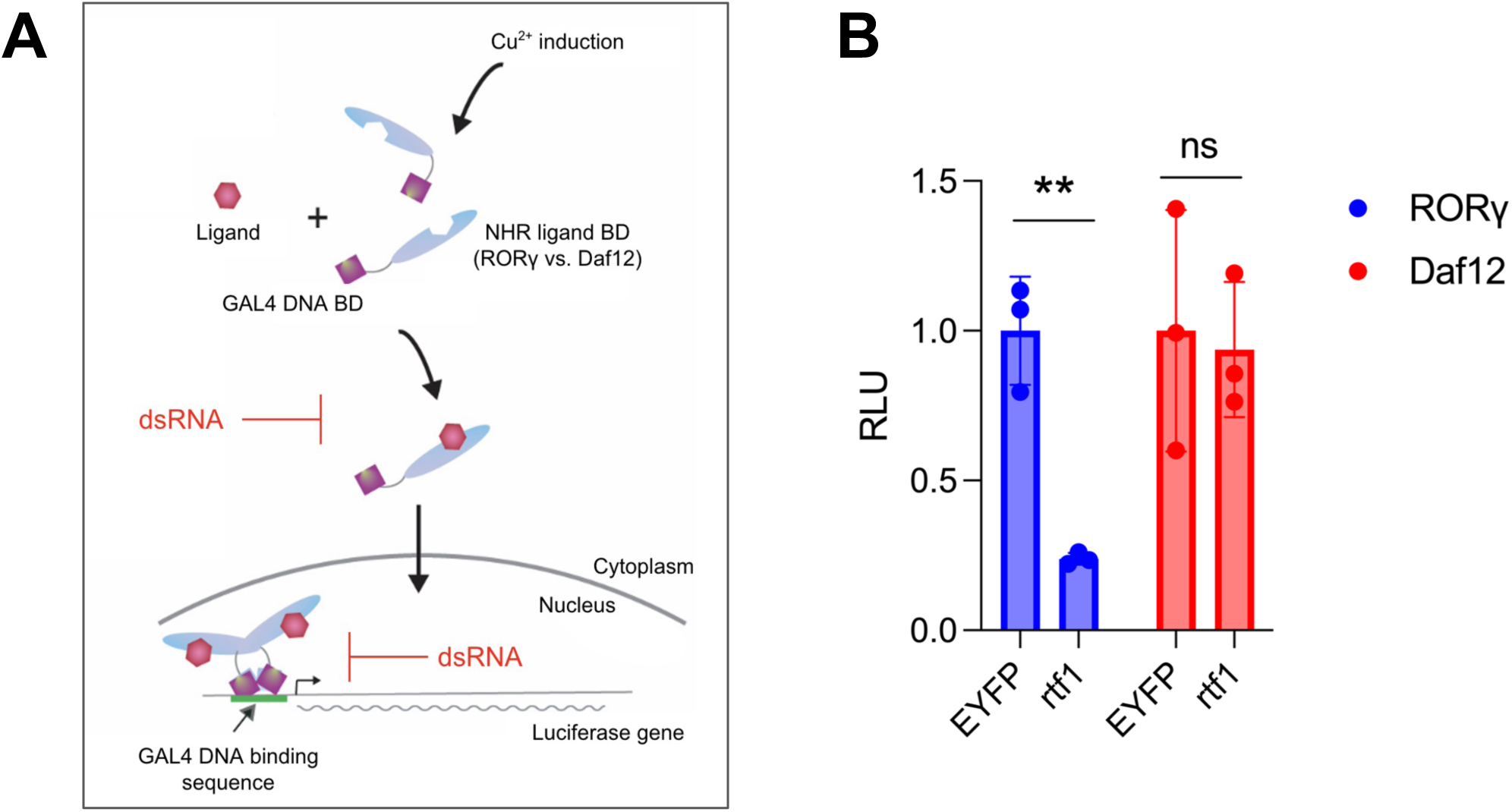
RTF1 regulates RORγ-driven transcription in the *Drosophila* S2 cell reporter system. **A**. Schematic depicting the S2 cell reporter system. A fusion protein containing GAL4 DNA-binding domain and NHR ligand binding domain from RORγ or Daf12 are expressed by pMT construct induced with Cu^2+^. After binding to the endogenous ligands, the fusion protein translocates into the nucleus and binds to the GAL4 DNA-binding sequence to drive the transcription of the luciferase gene. **B**. The reporter cells were treated with dsRNA targeting the *Drosophila rtf1* gene or EYFP as control. The activity of GAL4-RORγ or GAL4-Dap12-driven reporter transcription is measured by bioluminescence. The data represent analysis from 3 replicates (mean ± SD; unpaired *t*-test). \*\**P* < 0.01.

**Figure S2.**
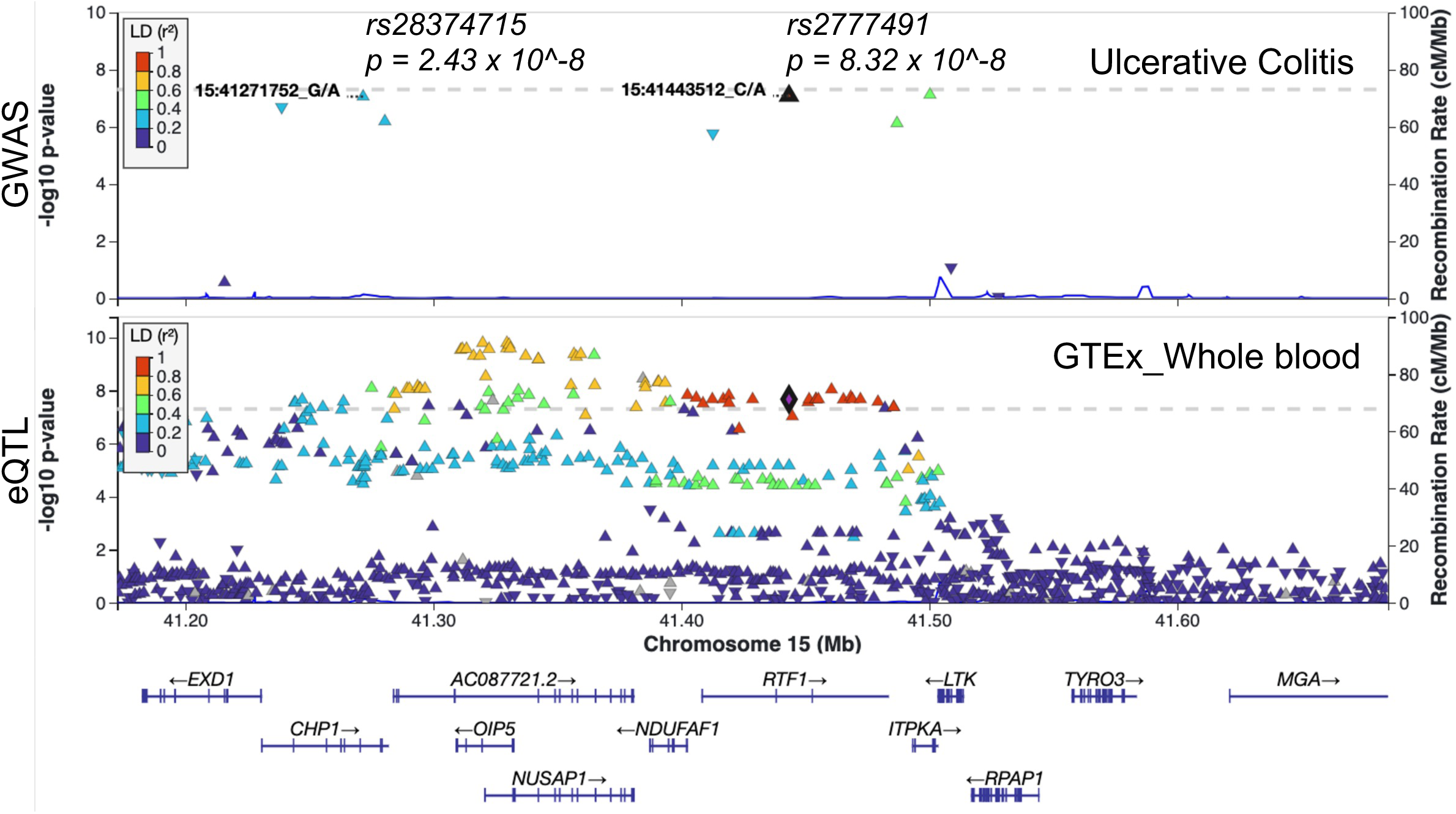
RTF1 genetic variants associated with a risk of ulcerative colitis. **A.** LocalZoom plot depicting regional association around *RTF1* represents the GWAS signal for ulcerative colitis, and the eQTL signal in whole blood from the GTEx v8 eQTL catalog The purple diamond indicates the variant rs2777491. The horizontal line indicates the genome-wide significance threshold (*P* < 5E-08).

**Figure S3.**
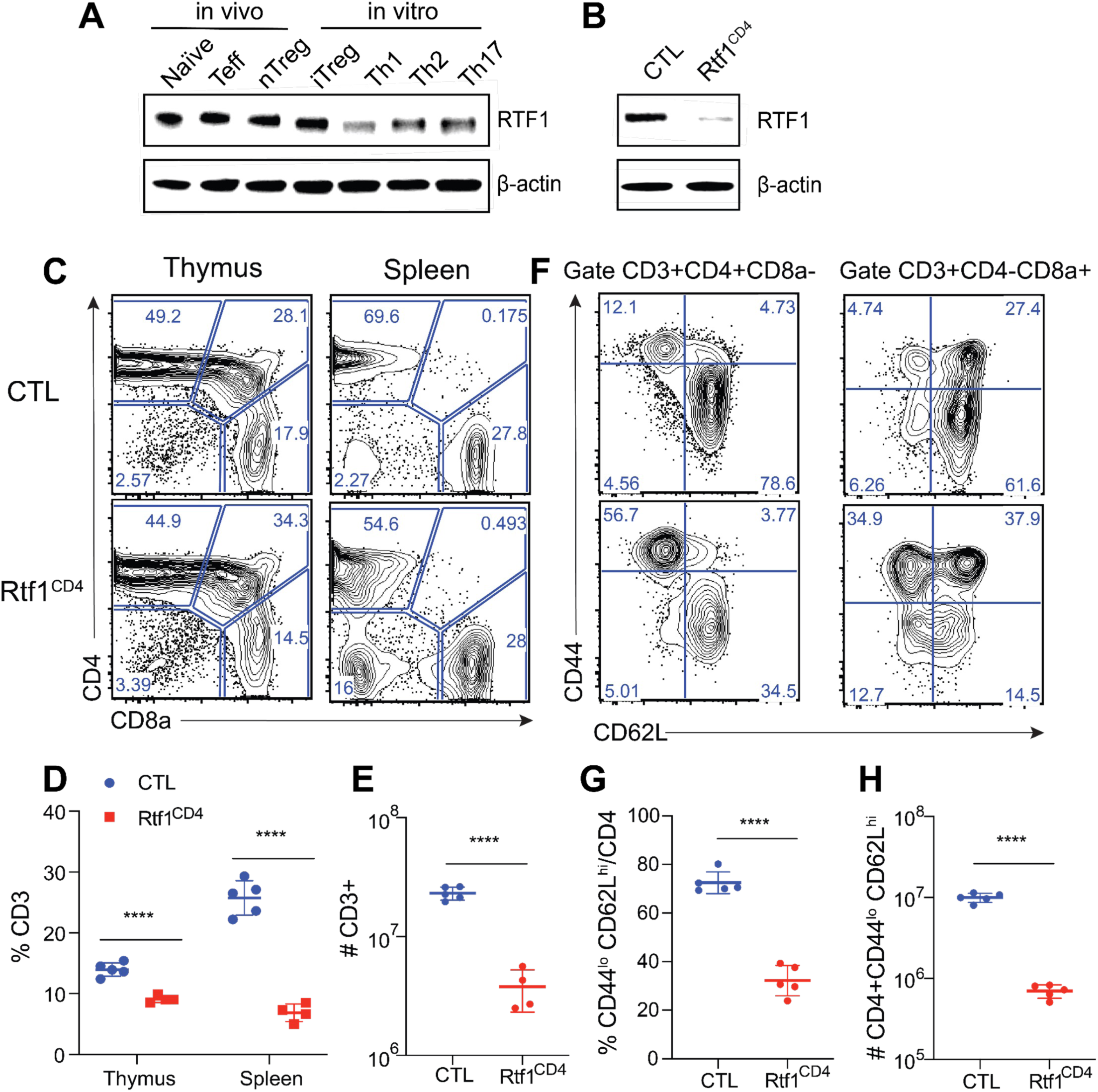
Impaired T cell development in Rtf1^CD4^ mice. **A**. Immunoblotting analysis of RTF1 expression in CD4^+^ naïve T, effector T (Teff), natural Treg (nTreg) cells isolated from B6 mice as well as in vitro-differentiated inducible Treg (iTreg), Th1, Th2, and Th17 cells. β-actin serves as a loading control. **B**. Immunoblotting analysis of RTF1 in total CD4^+^ T cells isolated from spleens and lymph nodes of Rtf1^fl/fl^ (CTL) and Cd4^Cre^Rtf1^fl/fl^ (Rtf1^CD4^) mice. **C**. Flow cytometric analyses of CD4 and CD8 expression on thymocytes and splenocytes from CTL and Rtf1^CD4^ mice. **D** and **E**. The percentage and total cell number of CD3^+^ T cells in the thymus and spleen of CTL and Rtf1^CD4^ mice. **F**. Flow cytometric analyses of CD62L and CD44 expression on the splenocytes of CTL and Rtf1^CD4^ mice. **G** and **H**. Percentage and absolute cell count of naïve CD4^+^ T cells in the spleens of CTL and Rtf1^CD4^ mice. Experiments were conducted on cells collected from 5 to 8 week-old mice. The data represent analysis from 4 to 5 mice per genotype (mean ± SD; unpaired *t*-test). \*\*\*\**P* < 0.0001.

**Figure S4.**
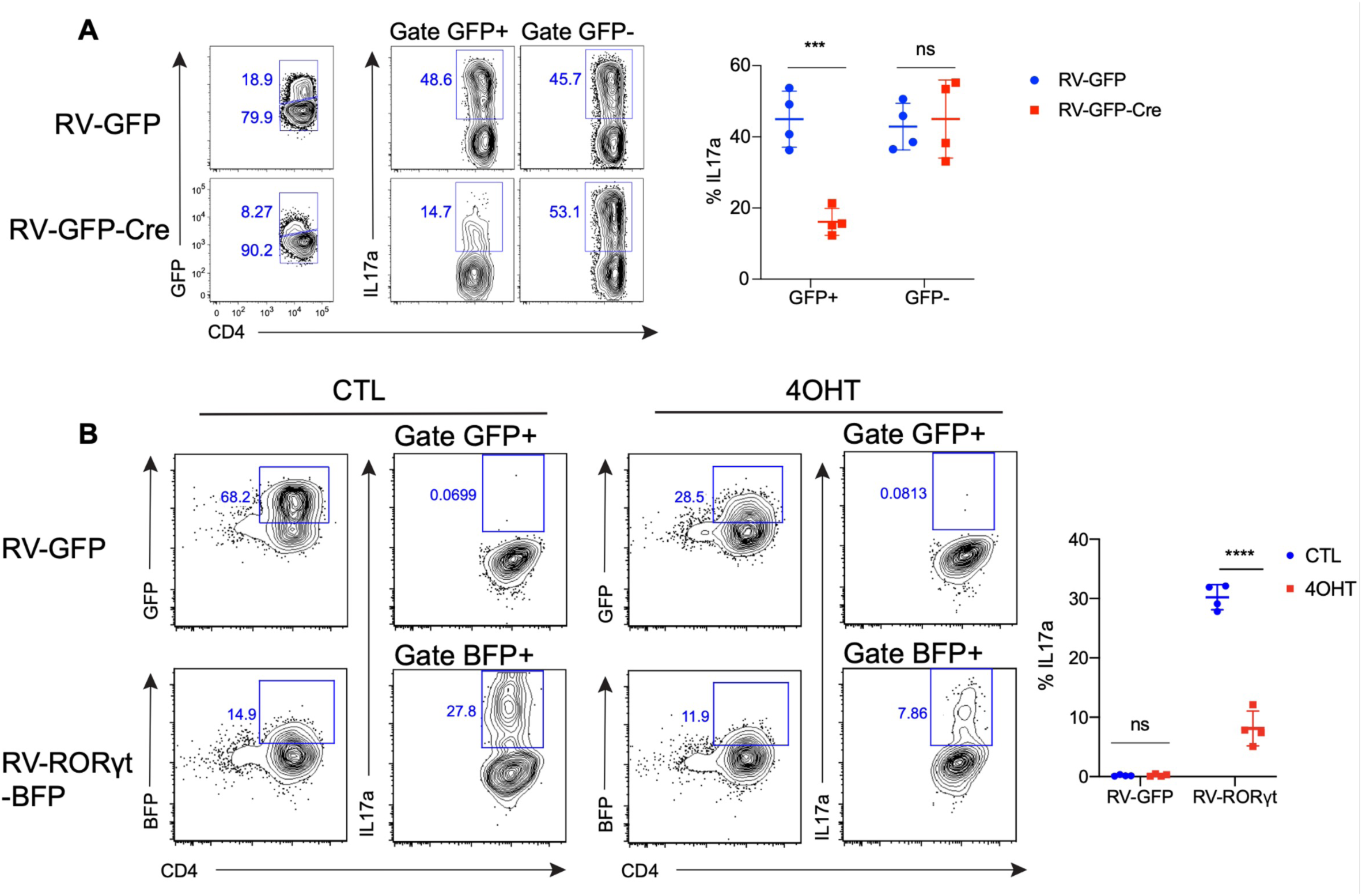
RTF1 is required for RORγt-dependent IL-17a expression. **A**. Flow cytometric and statistical analysis of in vitro cultured Th17 cells. Naïve CD4^+^ T cells from Rtf1^fl/fl^ mice were purified, cultured in vitro under Th0 (anti-CD3, anti-CD28, IL-2) conditions for 24 hours, transduced with retroviruses carrying GFP (RV-GFP) or GFP-Cre (RV-GFP-Cre), and polarized under Th17 (anti-CD3, anti-CD28, IL-6, TGFβ) conditions for an additional three days. **B**. Flow cytometric and statistical analysis of in vitro cultured Th0 cells. Naïve CD4^+^ T cells from UBC^Cre-ERT2^Rtf1^fl/fl^ (Rtf1^UBC^) mice were purified, cultured in vitro under Th0 (anti-CD3, anti-CD28, IL-2) conditions, and transduced with retroviruses carrying GFP (RV-GFP) or RORγt-BFP (RV-RORγt-BFP). Th0 cells were treated with DMSO (CTL) or 4-hydroxytamoxifen (4OHT) for an additional three days. Data represent three independently replicated samples (mean ± SD; unpaired *t*-test). ns = not significant. \*\*\**P* < 0.001, \*\*\*\**P* < 0.0001.

**Figure S5.**
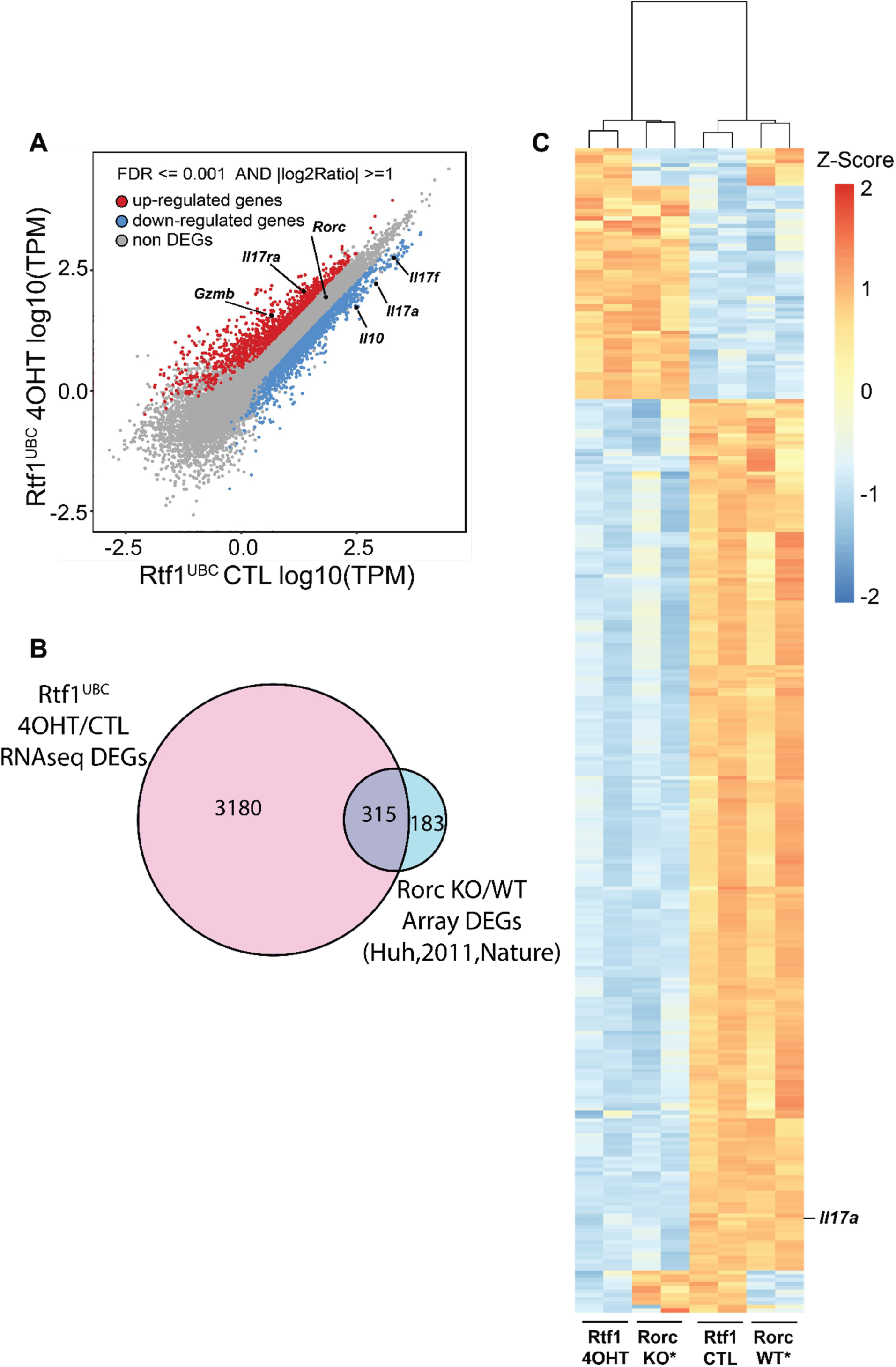
RTF1-dependent transcriptional program in Th17 cells. **A**. Scatter plot of Transcripts Per Million (TPM). RNA-seq comparing the gene expression profile of cultured Th17 cells from UBC^Cre-ERT2^Rtf1^fl/fl^ (Rtf1^UBC^) mice, treated with DMSO (CTL) or 4-hydroxytamoxifen (4OHT) for three days. **B**. Venn diagram of differentially expressed genes (DEGs) from the loss of Rtf1 (RNA-seq data from panel A) and the loss of Rorc (Microarray data from *Huh et al., 2011, Nature*) in cultured Th17 cells. **C**. Heatmap clustering of the 315 common DEGs from panel **B**.

**Figure S6.**
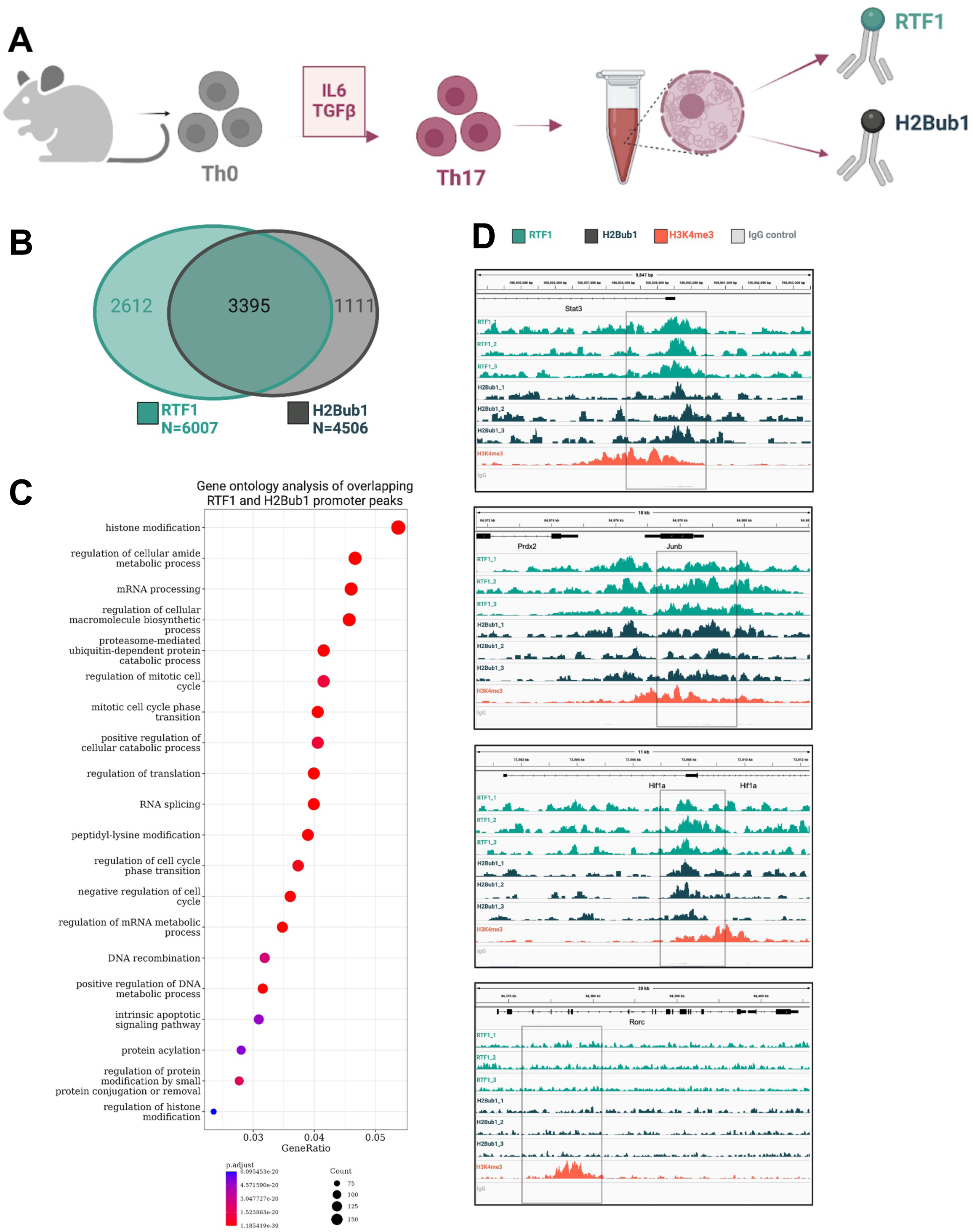
RTF1 and H2Bub1 share binding sites in Th17 cells. **A**. Schematic of the experimental setup for CUT&RUN. Th17 cells were differentiated in vitro with IL-6 and TGF-β for 3 days. Nuclei from Th17 cells were isolated for all downstream CUT&RUN experiments. CUT&RUN was performed for RTF1 and H2Bub1 as well as an H3K4me3 control and IgG control. **B**. Peak overlap for RTF1 and H2Bub1 whereby 3395/6007 (57%) of total RTF1 promoter peaks are shared with H2Bub1 and 3395/4506 (75%) of total H2Bub1 promoter peaks are shared with RTF1. **C**. Gene ontology analysis of 3395 shared peaks between RTF1 and H2Bub1. **D**. IGV panels of all three replicates of RTF1, H2Bub1, and control H3K4me3/IgG binding shown for *Stat3, Junb, Hif1a,* and *Rorc*; note the absence of RTF1 and H2Bub1 binding for *Rorc.* Annotated gene promoters shown in gray box Annotated gene promoters are boxed in gray.

